# A co-evolved peptide-GPCR system senses host entry to drive fungal infection

**DOI:** 10.1101/2025.09.26.678928

**Authors:** Gabriel Mendoza-Rojas, Philip Nakonz, Min Lu, Johannes Postma, Naomi Shtakser, Max Heinen, Manav Patel, Orlando Arguello-Miranda, Sonja Billerbeck, Florian Altegoer

## Abstract

A successful infection requires pathogens to recognize the specific host environment in order to reprogram their physiology accordingly. One major way in which eukaryotic cells sense their surroundings is via G-Protein Coupled Receptors (GPCRs), which share a seven-transmembrane architecture and G-protein-mediated downstream signaling. While mammalian GPCRs are well-characterized and represent important drug targets, their fungal counterparts remain poorly understood. In the corn pathogen *Ustilago maydis*, we now uncover a GPCR-based mechanism that allows the fungus to scout the host environment to sense whether it has entered into the plant tissue. During infection, the fungus secretes the protein Pit2, which is cleaved by host apoplastic-cysteine proteases, releasing a peptide ligand ‘hidden’ within Pit2. This ligand activates the fungal GPCR Gpe1 strongly promoting fungal proliferation after initial host penetration. Comparative analyses reveal conservation of the Gpe1/Pit2 system, with co-evolutionary signatures preserving receptor-ligand specificity. Furthermore, this GPCR system recognizing ‘hidden’ peptide ligands shows conceptual similarities to the fungal pheromone mating system, without sharing sequence similarity. Our findings reveal a co-evolved mechanism between fungus and host that encodes environmental context into a protein scaffold, establishing a novel paradigm for host-dependent signaling with implications for inter-organismic communication.

## Main

Interactions between pathogenic fungi and their hosts depend on the pathogens’ ability to perceive and respond to host-derived signals that facilitate infection. One well-characterized way of eukaryotic cells to sense the environment occurs via G-protein coupled receptors (GPCRs), which share a seven-transmembrane architecture and conserved intracellular signaling pathways. The human genome encodes over 800 GPCRs, which together represent targets of nearly 40% of approved drugs, highlighting their central roles in physiology and disease ^1^. Likewise, fungal genomes harbor numerous GPCRs, and emerging evidence indicates they contribute to colonization of humans and plants, although the repertoire of ligands and activation mechanisms remains poorly understood ^2–5^. The smut fungus *Ustilago maydis* is a pathogen of corn and a model organism for studying plant-pathogen interactions and fungal development. Like other smut fungi, *U. maydis* undergoes a complex life cycle that includes a yeast-like stage and infectious hypha ^6^. Transition to hyphal growth is driven by host-derived stimuli that induce expression of virulence-associated effector genes and control fungal proliferation - yet only a handful of such chemical signals have been characterized ^7^. Despite strong evidence for the importance of GPCR-mediated signaling ^6,8–11^, canonical GPCRs have not yet been investigated in *U. maydis*, leaving a significant gap in our understanding of fungal perception mechanisms in this pathogen.

### The GPCR Gpe1 is activated by a Pit2-derived peptide

To systematically explore GPCRs in *U. maydis*, we performed a genome-wide search for proteins containing the canonical seven-transmembrane (7TM) architecture using a structure-based approach utilizing Alphafold2 ^12^. This analysis identified 34 putative GPCRs (**Extended Data Tab.1**), of which only nine had been suggested by previous studies ^2,4,13^. Among the newly identified, one candidate was reported as crucial virulence factor 14, designated as **p**rotein **i**mportant for **t**umors 1 (Pit1) 14, which we renamed to **G**-protein-coupled receptor promoting fungal *in* ***p****lanta* **e**xpansion 1 (**Gpe1**).

Gpe1 is a 435-amino acid protein, featuring a short (35 aa) extracellular N-terminus and a longer (120 aa) unstructured intracellular C-terminus (**Fig.1a, Extended Data Fig.1**). Three extra- and intracellular loops connect the TMs, with the N-terminal 35 residues and the 38 residues of the second extracellular loop (ECL2) forming an extracellular extension (**Fig.1b**). ECL2 is also connected to TM3 via a disulfide bond formed by Cys108 and Cys178 (**Fig.1b**), a hallmark of many rhodopsin-class GPCRs ^15^. To explore a potential G-protein coupling, we used AlphaFold3-based ^16^ modeling to assess possible interactions between Gpe1 and the *U. maydis* Gα proteins. The analysis predicted an interaction with Gpa1 (confidence score: 0.58), while interactions with Gpa2 (0.23) and Gpa3 (0.21) were considered unlikely (**Extended Data Fig.1**). Based on the predicted interaction with Gpa1, we hypothesized that Gpe1 binds to this Gpa1.

**Figure 1.**
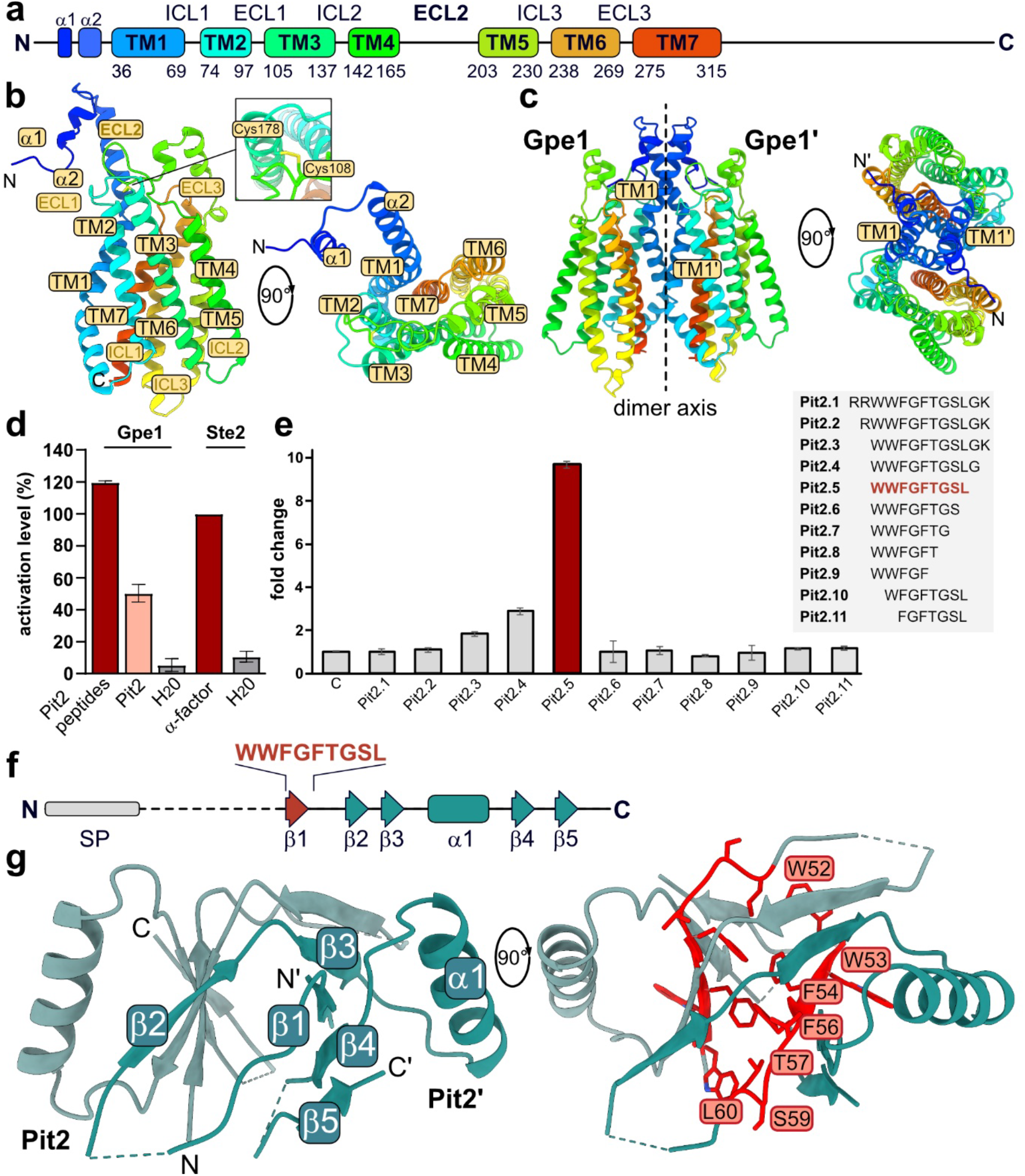
Structural and functional analysis of the GPCR Gpe1 and its Pit2 peptide ligand. **a.** Schematic representation of the domain architecture of Gpe1, indicating predicted transmembrane (TM) helices, intracellular (ICL) and extracellular loops (ECL), and N-/C-terminal regions. **b**. Side and top views of the predicted Gpe1 monomer structure with TM helices and loops labeled. A disulfide bond between Cys108 and Cys178 stabilizing the extracellular region is highlighted. Cartoon is colored in rainbow from N- (blue) to C-terminus (red). **c**. Dimeric arrangement of Gpe1 (Gpe1 and Gpe1’) shown along the membrane plane and rotated 90°, revealing the dimerization interface mostly involving TM1. **d**. Activation of Gpe1 by 100 µM full-length Pit2, 10 µM Pit2 peptides and *S. cerevisiae* Ste2 by 10 µM α-factor through fluorescent readout. Fluorescence levels are normalized to the Ste2 activation being 100 %. **e**. Fold-change activation of Gpe1 by a panel of self-secreted peptide variants (Pit2.1 to Pit2.11), with Pit2.5 (WWFGFTGSL) showing the strongest activation. Peptide sequences are listed on the right. **f**. Schematic of the Pit2 precursor protein, including its signal peptide (SP) and the position of the activating peptide (WWFGFTGSL) within the structural elements (β-strands and α-helices). **g**. Crystal structure of the Pit2 homodimer. Protomers are colored in cyan and light cyan. Secondary structure elements are labeled and the residues of the activating peptide buried in the dimer core highlighted in red.

To enable ligand identification, Gpe1 was expressed in an engineered *Saccharomyces cerevisiae* strain to test receptor activation through a fluorescent reporter ^17^. Western blot analysis confirmed expression of Gpe1 in *S. cerevisiae*, but predominantly detected a ∼110 kDa band, suggesting homodimerization of Gpe1 (**Extended Data Fig.2a**). This was further supported by structure prediction of the Gpe1 homodimer (confidence score: 0.57; **Extended Data Fig.2b**), indicating a dimer interface involving mostly the extracellular N-terminus and TM1 (**Fig.1c**).

The neighboring gene of *gpe1* encodes the secreted effector Pit2, which is cleaved by maize papain-like cysteine proteases (PLCPs) to release a protease-inhibitory peptide ^18,19^. Deletion of *gpe1* or *pit2* causes a phenotypically identically reduced virulence, although no functional link between both proteins has been established ^14^. We hypothesized that Pit2 might act as or contain the activating ligand of Gpe1 and purified recombinant Pit2 lacking its signal peptide (**Extended Data Fig.3a**). Purified Pit2 was applied to yeast expressing Gpe1, inducing a reporter response reaching ∼25% of the ScSte2 + α-pheromone control (**Fig.1d**). Because Pit2 is cleaved by maize PLCPs, we next tested whether cleavage enhances activity. Remarkably, tryptic digestion of Pit2 peptides - resulting in a similar cleavage pattern compared to PLCP treatment ^18,19^ - increased the reporter signal to ∼100% at tenfold lower peptide concentration (**Fig.1d**), demonstrating robust Gpe1 activation by processed Pit2 fragments in *S. cerevisiae*. Peptide fractionation and mass spectrometry identified an 11–amino acid Pit2 fragment (WWFGFTGSLGK) as a Gpe1 agonist (**Extended Data Fig.3b,c**). To delineate the core activation motif, we employed a yeast biosensor system featuring peptide secretion and monitoring of GPCR activation ^17^. This analysis mapped the core sequence to WWFGFTGSL (Pit2.5), triggering a nearly tenfold increase in fluorescence compared to the 11-amino acid sequence motif (**Fig.1e**). Intriguingly, the presence of an N-terminal RR-motif suppressed Gpe1 activation. Previous work had implicated a 14-residue, RR-containing Pit2 peptide (PID14) as PLCP-derived protease inhibitor ^18,19^. Consistently, the shorter WWFGFTGSL peptide also arises from PLCP cleavage but lacks protease-inhibitory activity ^18^. Together, these findings establish Gpe1 as a dedicated receptor for a Pit2-derived peptide, with activation strictly depending on host PLCP processing.

### Pit2 serves as scaffold for the Gpe1-activating peptide

Both, Gpe1 and Pit2 are crucial for plant infection and subject to tight regulation on the transcriptional and post-transcriptional level ^20,21^. Another layer of regulation seems to be contained in the Pit2 protein, which protects and hides the activating peptide prior to host-dependent activation. Consistent with that, Pit2 is secreted in full-length by *U. maydis* without evidence of endogenous cleavage ^20^. Size-exclusion chromatography of Pit2 further supported a stable protein scaffold, revealing a single monodisperse fraction (**Extended Data Fig.3a**). To understand the precise nature of this protein chassis, we crystallized Pit2 and solved its crystal structure at 2.3 Å resolution (**Extended Data Tab.2**). Residues 51 to 120 could be unambiguously placed in the electron density except for the N-terminus and flexible regions decorating the protein core. Pit2 is composed of five β-strands and one α-helix forming a tightly intertwined homodimer. The dimer interface buries the conserved hydrophobic WWFGFTGSL motif forming β1 with W52, W53, F54, and F56 contributing to the hydrophobic packing (**Fig.1g**). Strands β3-5 of one protomer and β2 from the neighboring Pit2 molecule complete the dimer interface (**Fig.1f,g; Extended Data Fig.4**). Our results indicate that Pit2 acts as a structural chassis containing the activating peptide in the homodimer interface, thereby preventing premature activation of Gpe1 prior to host-mediated proteolytic release of Pit2-derived peptides.

### Conserved residues at Gpe1 are required for Pit2 peptide binding

To investigate the structural basis of Gpe1 activation, we modeled Gpe1-Pit2 complexes using Alphafold3. In the predicted structure, the N-terminal tryptophan residues of the Pit2 peptide are buried top-down in a cavity formed by TM1 and ECL2 of Gpe1, with the C-terminus oriented extracellularly (**Fig.2a**). This arrangement delivers an explanation why N-terminal arginines block activation, whereas C-terminal extensions retain activity (**Fig.1e**).

**Figure 2.**
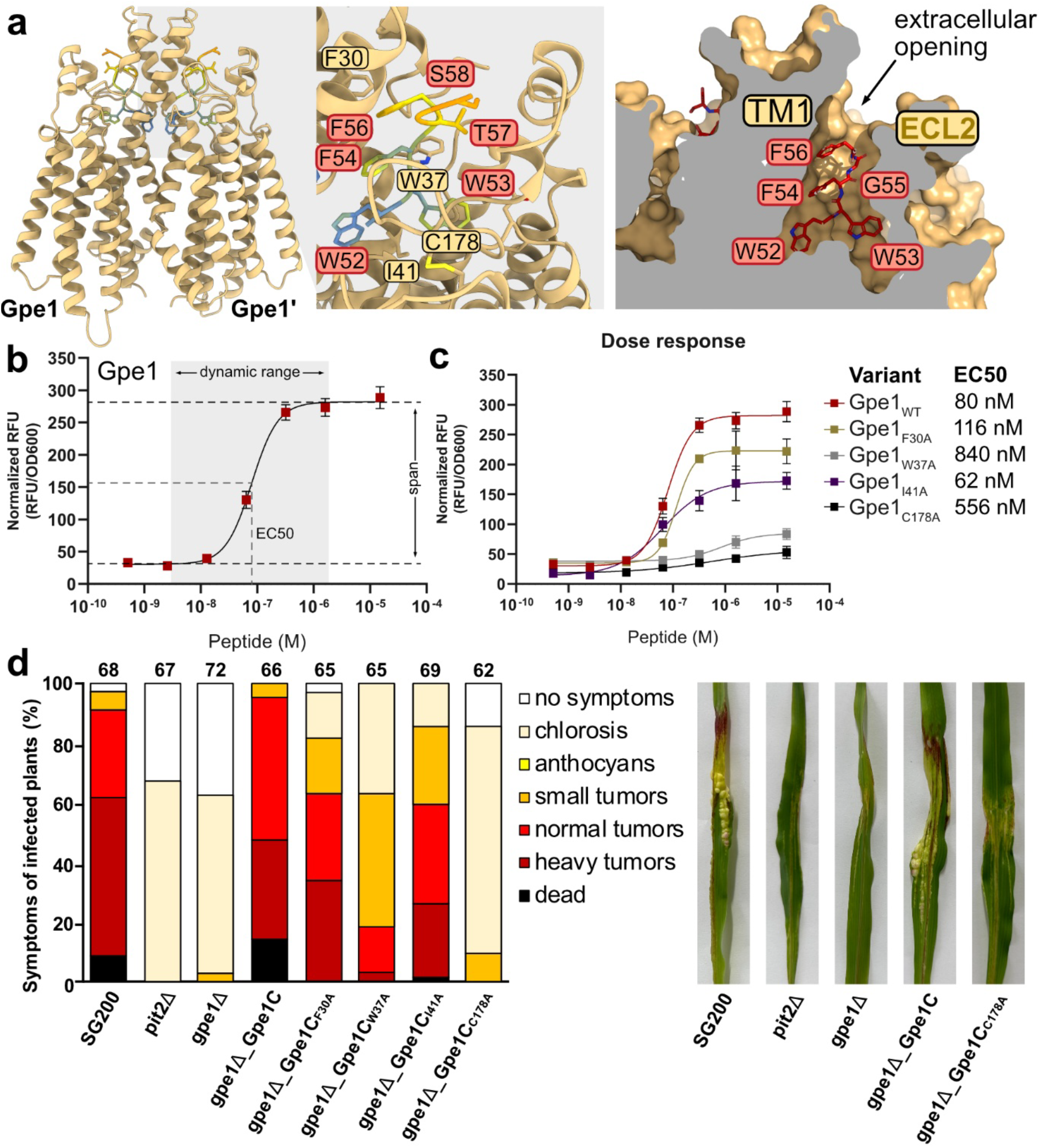
Structural determinants of Pit2 peptide binding and functional relevance of Gpe1. **a.** Predicted binding mode of the Pit2-derived peptide to Gpe1, highlighting residues at the interface (left). Close-up view shows aromatic and hydrophobic residues in TM1 and ECL2 (middle), with surface representation illustrating the extracellular opening of the binding pocket (right). **b**. Dose–response curve of wild-type Gpe1 activation by the Pit2 core peptide (WWFGFTGSL), showing normalized fluorescence reporter units (RFU) as a function of peptide concentration. Shaded area indicates dynamic range, with EC50 marked. **c**. Dose– response curves of Gpe1 variants mutated at predicted binding site residues, compared to wild-type Gpe1. Right: table summarizing EC50 values for the indicated variants. **d**. Maize seedling infection assays using *U. maydis* SG200 strains and derivatives thereof. Disease symptoms were rated 12 days after inoculation (dpi) and grouped into categories as shown in the legend. Numbers of infected plants are given above each column, representing three biological replicates. Images show representative infection symptoms.

Guided by these predictions, we generated Gpe1 variants with substitutions near the putative Pit2 binding site and expressed them in *S. cerevisiae* (**Fig.2a,b**). Dose-response analysis revealed an EC50 of 80 nM for wildtype Gpe1 (**Fig.2b,c**), consistent with values reported for fungal Ste2 pheromone peptide receptors ^17,22^. While some mutants (F30A, I41A and V166A/D167A) only moderately influenced EC50, dynamic range and maximum activation levels, variation of C177A and W37A strongly decreased EC50 and rendered Gpe1 almost non-responsive (**Fig.2c**).

Having established the *in vitro* importance of these residues, we next tested their functional relevance *in planta*. Deletion strains of *gpe1* were complemented with the different *gpe1* mutant alleles and maize infection assays were performed to assess virulence (**Fig.2d, Extended Data Tab.3**). As reported earlier, deletion of *gpe1* or *pit2* almost completely abolishes virulence preventing tumor formation (**Fig.2d**) ^14^. Complementation with the native *gpe1* and *gpe1*_*F30A*_ restored virulence to wildtype levels, while strains producing the Gpe1_I41A_ variant show a slight decrease and strains containing C177A and W37A a strong attenuation in virulence following the results obtained from *in vitro* experiments (**Fig.2c,d**). Collectively, our structure-guided mutational analysis confirmed the functional relevance of the Pit2 coordination by Gpe1 and revealed that disrupting this interaction strongly influences the virulence of *U. maydis*.

### Pit2-mediated activation of Gpe1 induces receptor endocytosis and promotes proliferation

Gpe1 and Pit2 are important regulators of fungal fitness during plant infection but the precise physiological impact is still unclear. To assess whether and how Gpe1 activation modulates fungal physiology, we constitutively expressed a green fluorescently tagged version of Gpe1 (Gpe1-G) in the haploid pathogenic *U. maydis* strain SG200.

Sporidial cells of the strain were subjected to a microfluidic microscopy setup that enables continuous incubation, live-cell imaging and controlled perfusion of the Pit2 peptide. In the absence of exogenous peptide, Gpe1-G displayed a plasma membrane-associated localization but also localized to vesicular structures (**Fig.3a,b**) supporting earlier observations^14^. Upon addition of activating Pit2 peptide, rapid internalization of Gpe1-G into the endomembrane system was observed, suggesting ligand-induced endocytosis and receptor recycling (**Fig.3a,b**). To further characterize the subcellular compartmentalization of internalized Gpe1, we performed co-localization experiments using organelle-specific dyes (**Fig.3c**). Vacuolar staining with FM4-64 showed that Gpe1-G mostly localizes to vacuoles, suggesting that the majority of internalized Gpe1 is trafficked to the vacuole for degradation (**Fig.3c**). These findings indicate that Pit2 stimulation triggers ligand-dependent endocytic trafficking of Gpe1, followed by receptor turnover via the vacuolar degradation pathway. Notably, we also observed that the Gpe1-G strain reacted with elevated growth rates. Upon Pit2 peptide exposure, individual fungal cells divided more rapidly, suggesting an accelerated cell cycle progression (**Extended Data Fig.5a,b**). More precisely, the time required to complete a cell division cycle (from one budding to the next budding event) decreased significantly from 226±40 min to 167±10 min upon treatment with Pit2 peptide (**Extended Data Tab.4**). In *U. maydis*, the infection process requires a complex transcriptional program inducing a G2 cell cycle arrest during mating, prioritizing polar growth during the early stage of plant colonization ^23^ (**Extended Data Fig.6**). The transcription factors bE/bW and Rbf1 control polar growth and cell cycle arrest ^23,24^ and the regulatory protein Clp1 has been implicated to play a decisive role in the proliferation vs. cell cycle arrest process ^25^. *In planta*, the cell cycle arrest is released and proliferation is prioritized again to enable massive fungal growth followed by gall formation. Clp1 is stabilized via the interaction with the central UPR regulator Cib1 to regulate pathogenic development at the onset of biotrophic growth *in planta* ^26^. Cib1 directly regulates expression of pathogenicity genes, including *gpe1* (*pit1*) and *pit2* ^20^. Our observation that activation of Gpe1 strongly promotes proliferation now suggests that activated Gpe1 acts via Clp1 (**Extended Data Fig.6**). To test this, we treated a *clp1* deletion strain with Pit2 peptides and monitored morphological changes. Strikingly, in a *clp1* deletion strain the Pit2 signal results in hyphal growth instead of proliferation (**Fig.3d**). This indicates that Clp1 is required to switch the cellular program from Rbf1-driven hyphal growth toward Cib1-dependent proliferation (**Extended Data Fig.6**). Likely, activation of Gpe1 supports stabilization of Clp1, potentially through a MAPK-dependent phosphorylation, although the precise mechanism is yet to be demonstrated. We therefore propose that Clp1 acts as a molecular switch, simultaneously repressing Rbf1 activity and stabilizing Cib1, thereby facilitating Gpe1-mediated proliferation during plant infection (**Extended Data Fig.6**).

**Figure 3.**
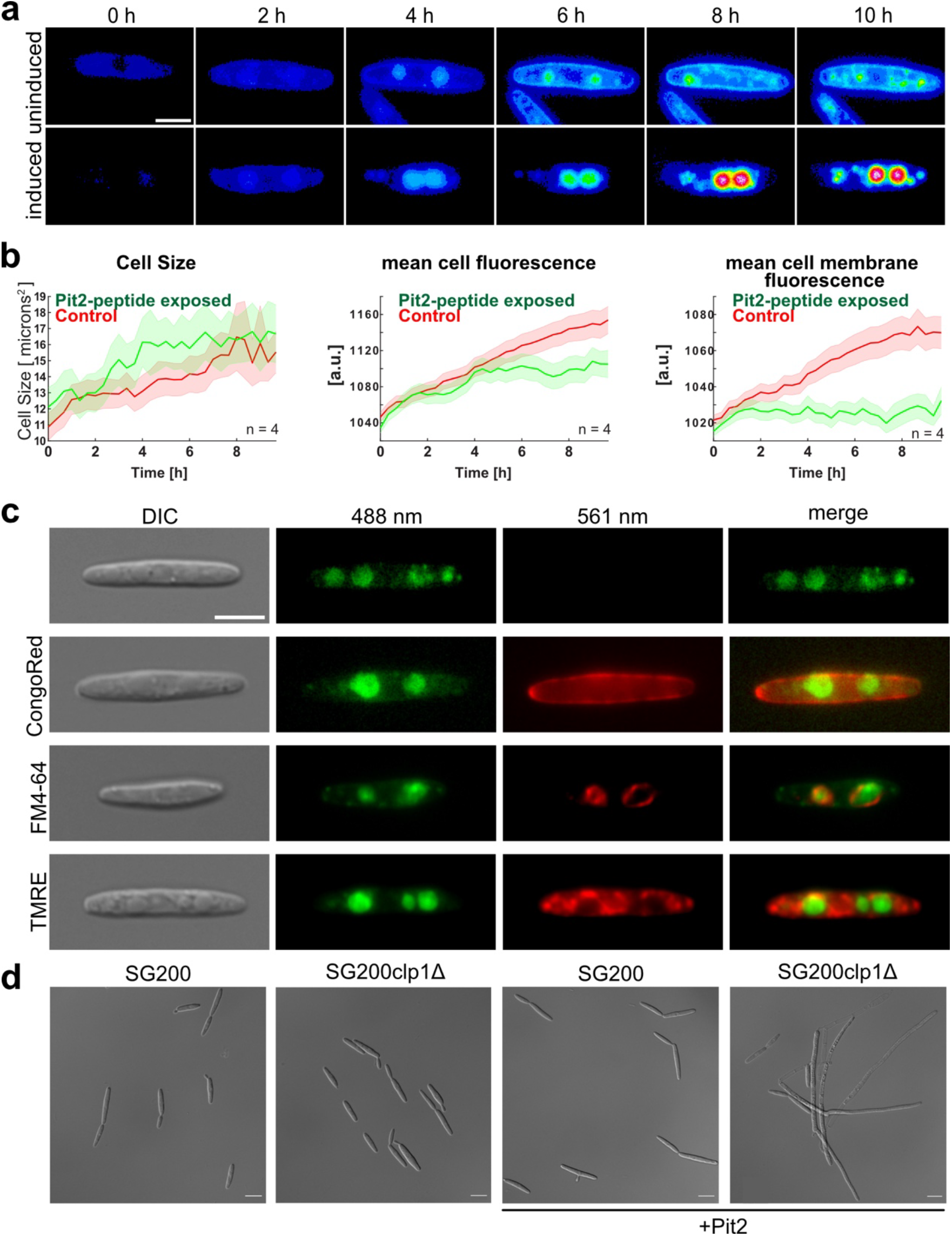
Gpe1 activation by Pit2 results in receptor internalization and promotes proliferation of *U. maydis*. **a.** Time-resolved imaging of Gpe1-G cells under uninduced or Pit2 peptide induced conditions using a microfluidic microscopy setup. Heat maps show fluorescence intensities at indicated time points. **b**. Quantitative analysis of cell size and mean cellular or membrane-associated fluorescence over time. Data represent mean ± s.e.m. from independent biological replicates (green, induced; red, uninduced). **c**. Subcellular localization of Gpe1-G in induced cells stained with Congo Red (fungal cell wall), FM4-64 (vacuolar membrane) and TMRE (mitochondrial membrane). Differential interference contrast (DIC), single-channel images with 488 nm or 561 nm excitation, and merged images are shown. Scale bars, 10 μm. **d**. DIC images of SG200 and SG200clp1Δ in absence and presence of 1 µM WWFGFTGSL Pit2 peptide. Peptide addition results in hyphal growth of SG200clp1Δ but not SG200.

### The Gpe1/Pit2 shows co-evolutionary signatures in smut fungi and similarities to the pheromone mating system

Gpe1 homologs are present in many smut fungi, whereas Pit2 is restricted to species closely related to *U. maydis* (**Fig.4a, Extended Data Tab.5**). Although Pit2 sequences display little overall conservation, the activating peptide residues are highly conserved (**Extended Data Fig.7**). To test cross-compatibility, we analyzed Pit2 homologs from *S. reilianum* (SrPit2) and *U. hordei* (UhPit2). Peptides corresponding to the *U. maydis* activating motif were assayed with the respective Gpe1 homologs in *S. cerevisiae*. SrPit2 activated UmGpe1 to levels comparable with UmPit2, whereas UhPit2 failed to do so (**Fig.4b**). Conversely, UmPit2 and SrPit2 only weakly stimulated UhGpe1 (30% and 45% activity, respectively; **Fig.4b**). These reciprocal activities suggest co-evolution of Gpe1 and Pit2, where minor sequence differences reduce cross-activation (**Fig.4b**). We conclude that the Gpe1/Pit2 module is broadly present in smut fungi and relies on host PLCP processing to generate peptides that both activate Gpe1 and inhibit PLCPs. This is further supported by the broad relevance of plant proteases for development and plant immunity. Several hundred proteases regulate central developmental programs in plants ^27^ and more recent studies highlight their central role in immunity ^28,29^. PLCPs are broadly conserved in plants and often strongly induced in different plant tissues upon pathogen infections and stresses ^29^. The Gpe1/Pit2 system therefore hijacks a central and indispensable aspect of plant immunity, potentially balancing immune suppression with fungal proliferation *in planta* (**Fig.4c**).

**Figure 4.**
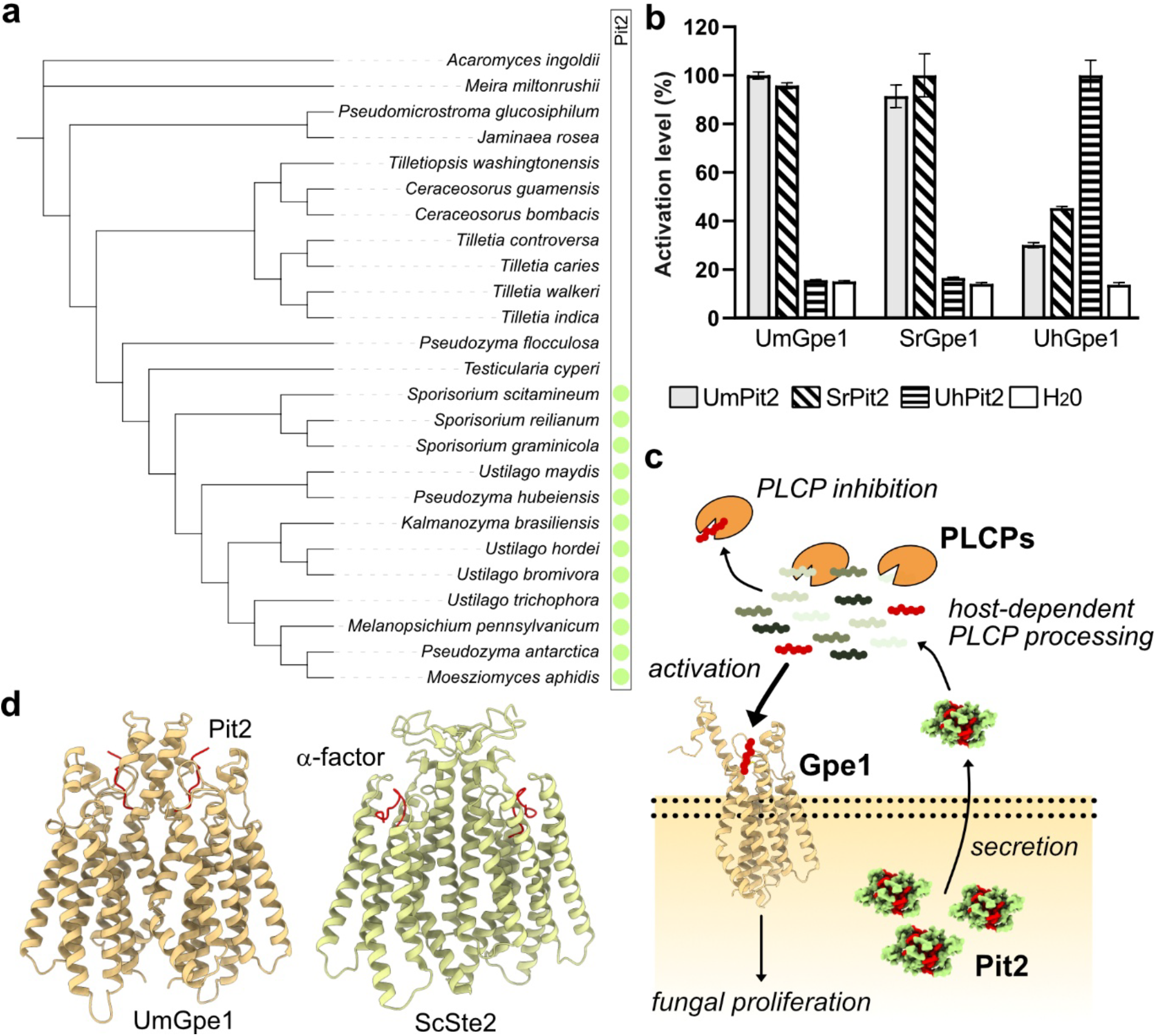
Role and co-evolution of the Pit2–Gpe1 system in smut fungi and shared features with the fungal pheromone mating system. **A.** Phylogenetic distribution of Gpe1 and Pit2 across smut fungi. Presence of Pit2 is indicated by a green colored circle. **b**. Activation of Gpe1 homologs from *Ustilago maydis* (UmGpe1), *Sporisorium reilianum* (SrGpe1), and *Ustilago hordei* (UhGpe1) by corresponding Pit2-derived peptides measured in *S. cerevisiae* biosensor assays. Bars show mean reporter activity ± s.d. **c**. Schematic model of Gpe1-Pit2 function. Secreted Pit2 is processed by host papain-like cysteine proteases (PLCPs), generating peptides that activate Gpe1 and inhibit the PLCPs. Gpe1 activation triggers fungal proliferation. **d**. Structural comparison of Pit2-bound UmGpe1 (left) with α-factor bound *S. cerevisiae* Ste2 (right), highlighting overall structural similarity of the two GPCRs

Despite lacking sequence homology, the Gpe1/Pit2 signaling system shares key features with the fungal pheromone mating system. Likewise, both receptors form homodimers ^30,31^, recognize short hydrophobic peptides ^32^ and share overall structural similarity (**Fig.4d**). However, unlike mating pheromones that are secreted and processed by fungal enzymes ^33^, Pit2 requires cleavage by host-derived PLCPs to release the ‘hidden’ activating peptide ^18,19^. In conclusion, our work identified a host-dependent peptide–GPCR signaling system in smut fungi, mediating fungal recognition of the host-specific plant environment and expanding the repertoire of peptide-based communication systems in fungi. Hence, similar “host-gated” signaling modules may be more widespread in pathogenic and symbiotic fungi than previously recognized. From an evolutionary perspective, the parallels between Pit2-derived signaling and fungal pheromone systems raise the possibility that fungi have repeatedly co-opted short, protease-processed peptides as ligands for GPCRs, tailoring these systems to different ecological contexts. Exploring peptide-GPCR pairs across fungal lineages may therefore uncover a hidden communication network that integrates fungal life stages within microbial communities and host environments.

## Acknowledgements

We thank Kai Heimel, Alexej Kedrov, Michael Feldbrügge and Kerstin Schipper for critical reading and fruitful discussions on the manuscript. We acknowledge support of peptide HPLC-separation by Jens Reiners and the Molecular Proteomics Laboratory (MPL) for mass spectrometry analysis.

## Author contributions

Conceptualization: F.A. Methodology: G.M., O.A.M., S.B. & F.A. Investigation: G.M., P.N., M.L., H.P., N.S. & M.P. Validation: G.M. & P.N. Formal analysis: G.M. & O.A.M Resources: O.A.M, S.B., F.A. Writing Original Draft: F.A. & G.M. Writing Review & Editing: all authors. Visualization: G.M., P.N., M.L., O.A.M & F.A. Supervision: F.A.

## Data and Code availability

Coordinates and structure factors have been deposited within the protein data bank (PDB) under accession codes: 9RXK. The authors declare that all other data supporting the findings of this study are available within the article and its supplementary information files. Original Python and MATLAB code used for image analysis and single cell tracking are available at https://github.com/MirandaLab/Peptide-GPCR_system_Single_Cell_quantifications

## Methods

### Accession numbers

*U. maydis, S. reilianum* and *U. hordei* Gpe1-Pit2 sequences were obtained from FungiDB (https://fungidb.org/fungidb/app/): UmGpe1 (UMAG_01374), UmPit2 (UMAG_01375), SrGpe1 (sr10528), SrPit2 (sr10529), UhGpe1 (UHOR_02063) and UhPit2 (UHOR_02064).

### Strains and growth conditions

All strains used and generated in this study are listed in **Extended Data Table 6**. The *Escherichia coli* strain TOP10 (Thermo Fisher Scientific) was used for cloning purposes. The *E. coli* strain SHuffle® (DE3) (Novagen) was used to express all produced proteins in this study. *E. coli* strains were grown in dYT medium (1.6 % (w/v) Bacto-Tryptone, 1 % (w/v) yeast extract and 0.5 % (w/v) sodium chloride) or Luria Broth (LB) media, supplemented with chloramphenicol (50 µg/mL) or ampicillin (100 µg/mL) under constant shaking (200 RPM) in a temperature-controlled incubator at 37 °C or on dYT-agar at 37 °C. *Zea mays* cv. Early Golden Bantam (EGB, Urban Farmer, Westfield, IN, USA) was used for infection assays with *Ustilago maydis* and grown in a temperature-controlled greenhouse (light and dark cycles of 14 hours at 28 °C and 10 hours at 20 °C, respectively). *U. maydis* strains were grown in YEPSlight medium (1 % (w/v) yeast extract, 0.4 % (w/v) peptone and 0.4% (w/v) sucrose) at 28 °C with baffled flasks under constant shaking at 250 rpm or on potato dextrose-agar at 28 °C. *Saccharomyces cerevisiae* strain *JTy014* was grown in Synthetic Complete (SC) media (all amino acids and 2% (w/v) glucose, pH 6.5) or on potato dextrose-agar at 28 °C. *S. cerevisiae* transformants which express GPCRs were grown in Synthetic dropout media (SD) media (SC media lacking uracil), Yeast Nitrogen Base (YNB) media (plus 0.01 % (w/v) methionine, 0.008 % (w/v) histidine, 0.04 % (w/v) leucine and 2% (w/v) glucose, pH 6.5) for fluorescence measurements or SD-agar at 28 °C. *S. cerevisiae* transformants with peptide secretion vectors were grown in synthetic dropout media (SC media lacking uracil and histidine) media with 1% sucrose instead of glucose (1% galactose was added to induce peptide expression).

### DNA amplification and molecular cloning

All plasmids and oligonucleotides used in this study are listed **Extended Data Table 6**. The open reading frames of all effectors cloned in this study were amplified from the genomic DNA of *U. maydis, S. reilianum* and *U. hordei*. All genes were amplified without their predicted signal peptide (SignalP 5.0) (Almagro Armenteros et al., 2019) employing specific oligonucleotides. A standard PCR protocol with Q5® High Fidelity DNA polymerase (New England Biolabs) and primer-specific annealing temperatures was used for the DNA amplification.

For the plasmid constructions, standard molecular cloning strategies and techniques were applied. For GPCR expression in *S. cerevisiae*, the ScSte2 expression vector pRS416 ^17^ was digested with *SpeI* and *XhoI* to remove the ScSte2 gene. The pure backbone vector was used for classical restriction-ligation (T4 ligase) to clone the GPCR genes inside using unique restriction sites (*SpeI* and *XhoI*) added in the PCR gene amplification. For plasmid amplification, recombinant plasmids were transformed in chemically competent *E. coli* Top10 cells. For site directed mutagenesis in the *gpe1* wild-type gene, whole-plasmid amplifications were performed using a standard PCR protocol with Q5® High Fidelity DNA polymerase (New England Biolabs), primers carrying a different codon for alanine instead the targeted amino acid, and as a DNA template the UmGpe1 expression vector (pIL0186).

Multiple plasmids from the MoClo yeast toolkit ^34^ were used to construct peptide secretion vectors. The GFP-dropout fragment was PCR-amplified to add *BsmBI* sites at both ends, followed by digestion with *BsmBI* (NEB). The secretion cassette of α-factor was obtained by amplifying pSB21 using primer ML215 and ML216. The YTK plasmids and pre-pro region PCR products were first assembled using Golden Gate system, then ligated with BsmBI-digested GFP-dropout product to form the final plasmid. All Pit2 variants were synthesized as single-stranded oligonucleotides and double-stranded DNA was generated either by PCR using a reverse primer SB320 or by annealing complementary primers. The resulting DNA fragments were inserted into the secretion vector using Golden Gate assembly. A GAL1 promoter was used to drive peptide expression.

For cloning of protein expression constructs, the pEM-GB1 vector was used for, encoding the solubility tag GB1, a hexahistidine (6His) tag and a *Tobacco etch virus* protease (TEV) recognition site. Plasmid assembly was performed as described previously ^35^. All plasmids were verified by Sanger sequencing.

### Transformation and generation of *S. cerevisiae* strains

A main culture of the respective *S. cerevisiae* ancestor strain was inoculated with an overnight preculture at a starting OD600 of 0.15 and incubated at 28 °C for 3-5 h. Subsequently, the main culture was harvested and washed with sterile water. Washed cells were resuspended in 1 mL of 100 mM of lithium acetate. Cells were transferred to 1.5 mL centrifuge tubes, harvested again and resuspended now in 0.1 mL of 100 mM of lithium acetate per transformant (0.5 mL for 5 transformants). For each transformant, 0.1 mL of resuspended cells were transfer in a new 1.5 mL centrifuge tubes, centrifuged and the following solutions were added in this order: 240 μL of 50 % (w/v) PEG 3350, 36 μL of 1M lithium acetate, 20 μL of boiled carrier DNA (salmon testes DNA, 5 mg/mL), 2 μL (∼2 μg) of plasmid (except for the negative control which sterile water was used instead) and necessary sterile water until a final volume of 360 μL. The transformation reaction was vortexed vigorously until the cell pellet was completely resuspended. Then, two incubation steps were performed without shaking: 30 °C for 30 min and 42 °C for 35 min. After a quick centrifugation the supernatant was discarded, and the cells were resuspended in 400 μL of sterile water smoothly using the micropipette. Onto two selection plates (SD-agar) the cells were streaked, one plate with 100 μL and the second one with 300 μL. The plates were incubated at 28 °C for 2-3 days.

### Western Blotting

*S. cerevisiae* strains were grown in 50ml SC-Ura medium to an OD_600_ of 1. Cells were harvested at 500 x *g* for 2 minutes and resuspended in 500 µl cold 1x PBS. Glass beads were added before processing in a Mixer Mill MM 400 (Retsch GmbH, Haan, Germany) at 30 Hz for 5 minutes. Cell debris was harvested at 300 x *g* for 5 minutes at 4°C. The supernatant was then centrifuged at 16000 x *g* for 45 minutes at 4°C and the membrane fraction containing pellets were resuspended in 50µl cold 1x PBS. 20 µl of the sample was mixed with 8 µl 5x Laemmli buffer (62.5 mM Tris-HCl, 2% SDS, 10% glycerol, 0.01% bromophenol blue, 5% β-mercaptoethanol, pH 6.8) and incubated at 50°C for 10 minutes.

10 µl of the sample was loaded onto a 12% SDS-PAGE gel and run at 100 V for 10 minutes followed by 180 V for 40 minutes. Blotting is performed using PVDF membrane at 45 mA for 1 hour, with subsequent blocking in TBST (20 mM Tris-HCl, 150 mM NaCl, 0.1% Tween-20, pH 7.4–7.6) + 3% milk powder overnight. The membrane was then incubated with Mouse Anti-HA antibody (Roche Diagnostics GmbH, Mannheim, Germany) for 2.5 h, washed three times with TBST and then incubated with Anti-Mouse IgG-HRP antibody (Promega Corporation, Madison, WI, USA) for 1.5 h. Following three washes with TBS (20 mM Tris-HCl, 150 mM NaCl, pH 7.4– 7.6), the membrane was incubated with ECL Prime (Cytiva, Marlborough, MA, USA) for 1 minute. Detection was performed using a ImageQuant LAS4000 (Cytiva, Marlborough, MA, USA).

### Protein production and purification

Plasmids were transformed into *E. coli* strain BL21 (DE3) (Novagen). Expression test conditions were induced in dYT median supplemented with the required antibiotic and lactose 1% for 20 h at 28 °C, IPTG 0.5 mM for 3 h at 37 °C or IPTG mM for 20 h at 20 °C, respectively at 200 RPM. After protein expression at the best condition, the cells were harvest (4 000 RPM, 15 min, 4 °C), washed, resuspended in lysis buffer (20 mM HEPES pH 8, 20 mM KCl, 250 mM NaCl and 40 mM imidazole) and lysate using a microfluidizer (M110-L, Microfluidics). All *E. coli* debris were collected by centrifugation (20 000 RPM, 15 min, 4 °C) and membrane free supernatant was loaded onto Ni-NTA FF-His Trap columns (GE Healthcare) for affinity purification. The elute protein in high-imidazole buffer (20 mM HEPES pH 8, 20 mM KCl, 250 mM NaCl and 500 mM imidazole) was subjected to a buffer exchange with no imidazole buffer (20 mM HEPES pH 7.5, 20 mM KCl, 200 mM NaCl) using Amicon Ultra-10K centrifugal filters (Milipore), added TEV protease in a protease:protein ratio (1:100 OD280) and incubated in a rotation well at 4 °C overnight (∼16 h). The cleaved protein was subjected to a reverse affinity chromatography to remove the hexahistidine tag fused with GB1 and uncleaved protein. The flowthrough was collected, concentrated to 1-2 mL using Amicon Ultra-10K centrifugal filters (Milipore) and subjected to size-exclusion chromatography (SEC) using a Superdex S200 Increase 16/600 column. The peak fractions were analyzed using a standard SDS-PAGE protocol, pooled, and concentrated with Amicon Ultra-10K centrifugal filters. Protein aliquots were flash-frozen in liquid nitrogen and then stored at −80°C.

### Protein crystallization

Crystallization was performed by the hanging-drop method at room temperature in 2 μL drops (1:1, protein:precipitation solution). Hexahistidine tag fused with GB1-UmPit2 crystallized at 20 mg/mL within 2 weeks in in 0.1 M Bicine and 1.6 M Ammonium sulfate at pH 9. Crystals were frozen flash-frozen in liquid nitrogen using a cryo-solution (mother solution supplemented with 20% (v/v) glycerol.

### Structure analysis by X-ray crystallography

Data were collected under cryogenic conditions at the European Synchrotron Radiation Facility (ESRF) beamline ID23-2 ^36^. The data were integrated and scaled using XDS and merged with XSCALE ^37^. The structure of Pit2 was determined by molecular replacement using Phenix ^38^ integrated Phaser-MR and 5BMH as search model. The structure was manually built in Coot ^39^ and refined in Phenix^38^. All residues were found within the preferred and additionally allowed regions of the Ramachandran plot. Detailed data collection and refinement statistics are listed in **Extended Data Table 2**.

### Structure prediction

Protein structures and interactions were predicted by AlphaFold 3 ^16^ and visualized using PyMOL (https://pymol.org/) or ChimeraX-1.9 ^40^.

### Plant infection assays

*U. maydis* strains were grown in 50 mL YEPS_light_ medium from an overnight pre-culture to an OD600 of 1. Cells were harvest by centrifugation (3 500 RPM, 5 min, room temperature), and resuspended in double-sterilized H_2_O to reach a final OD600 of 3. For the infection of maize plants, 250 - 500 μL of culture was injected into 7-day-old maize seedlings using a syringe. Disease symptoms were scored after 6-and 12-days post-infection and based on three biological replicates per *U. maydis* strain. Disease symptom scoring data are listed in **Extended Data Table 3**.

### GPCR activation assay in *S. cerevisiae*

The activation assays for each GPCR were performed as previously described with slight modifications ^17^. The peptides from Pit2 variants were obtained through trypsin digestion (1:20, trypsin:protein ratio) 37°C overnight (∼12-15 h) in TE buffer (20 mM Tris-HCl pH 8, 2 mM EDTA) or synthesized (Genscript Biotech). Experiments were performed in 96-well microtiter plates using a microplate reader (Tecan) and OD_600_ and red fluorescence were measured after 8h. Experiments were run in triplicate.

For dose response curves, 9 different concentrations (15 μM and 6 five-fold dilutions starting at 8 μM peptide, sterile water was used as no peptide control) of the specific synthetic peptide were used. Experiments were performed in 96-well microtiter plates using a microplate reader (Tecan) and OD_600_ and red fluorescence were measured after 20h. A four-parameter non-linear regression model (curve fit) was used using Prism (GraphPad): [Agonist] vs. response (variable slope). After analysis, EC_50_, basal activation, maximal activation and the Hill coefficient was extracted as previously described^17^.

The peptide secretion and GPCR expression strains were cultivated in SD media overnight, and diluted 1:1 with sterile water. Ten microliters cells were seeded in 200 μL total volume in 96-well plates and cultured at 30 °C with 800 rpm shaking. The fluorescence intensity was measured at 20h using microplate reader (Tecan) with excitation at 588 nm and emission at 620 nm. The fold change was calculated by dividing OD_600_-normalized fluorescence values of Pit2 variants by that of the control (cells containing empty plasmid), with the control value set to 1.

### Reversed-phase Pit2 peptides isolation by HPLC

The mixture of tryptic-digested UmPit2 peptides were subjected to separation with a binary gradient of from 5 % to 95 % acetonitrile with a flowrate of 2 mL/min on a HPLC system (Agilent) for 60 min operated in a C-18 reverse phase column. The injection point was after 2 min and while the separation process was running, 2 mL sample was collected until fraction number 50. Fractions which contain peaks at 205 nm or 280 nm, were incubated at 98 °C with the lid open under a fume hood until the samples are dried. Dried peptides were resuspended in 100 μL PBS buffer (20 mM HEPES pH 7.4, 20 mM KCl, 100 mM NaCl and 10 mM Na_2_HPO_4_). 15 μL of each selected fraction was tested for GPCR activation assay in *S. cerevisiae* as described before.

### Fluorescence Microscopy

Laser-based epifluorescence microscopy was performed on a Zeiss Axio Observer.Z1 equipped with Prime BSI Express SCMOS camera (Visitron Systems, Puchheim, Germany). Excitation was provided by a VS-LMS4 Laser-Merge-System (Visitron Systems, Puchheim, Germany) that combines solid-state lasers targeting GFP (488 nm at 50 or 100 mW) and RFP (561 nm at 50 or 150 mW). An emission filter for Gfp of 520 nm and for Rfp of 631 nm was used. Image acquisition was performed using VisiView (Visitron Systems, Puchheim, Germany).

Cells were grown in 10 ml cultures with 1µM WWFGFTGSL peptide to an OD_600_ of 0.4 prior to fluorescence imaging. All staining was imaged using RFP excitation. Congo Red staining involved labeling 2 µl of cells with 3 mM Congo Red directly on the slide. FM4-64 staining required incubating 1 ml cell samples with 8 µM FM4-64 at room temperature over 15 minutes, followed by one wash with 1x PBS (137 mM NaCl, 2.7 mM KCl, 10 mM Na_2_HPO_4,_ 1.8 mM KH_2_PO_4_). TMRE staining involved incubating 1.5 ml samples with 10 µM TMRE at 28 °C over 20 minutes, followed by two washes with 1x PBS. Image processing was carried out using Fiji ImageJ ^41^.

### Microfluidics Microscopy

Pictures were taken with an openFrame Microscopy System (Cairn GmbH, Ettlingen, Germany) equipped with the SCMOS sensor KINTEIX-22MM-M-C (Teledyne Photometrics, Tucson, AZ, USA) and an LDI-7 solid-state laser (89north, Williston, VT, USA) as illuminator. GFP imaging utilized 470 nm excitation and a 510 nm emission filter. Image acquisition was performed using Micro-manager 2.0 ^42^. Signal visualization was enhanced by applying the 16 colors lookup table to the GFP signal. Image processing was carried out using Fiji ImageJ ^41^.

Cells were grown in 10 ml cultures to an OD_600_ of 0.2. Following dilution with H2O to an OD_600_ of 0.05, 200 µl of these cells was added to a µ-Slide I^0.2^ Luer Uncoated (Ibidi, Mertinsried, Germany). NM medium containing 1 µM WWFGFTGSL peptide was continuously flushed over the cells throughout the entire duration using an EasyFlow Pump System (Fluigent, Paris, France) at a flow rate of 0.4 µl/min. Images were captured every 20 minutes over a 10-hour period.

### Microscopy data evaluation and single cell tracking

Single fungal cells in microfluidic devices were tracked with the FIEST algorithm, which uses custom-trained cellpose models and generative frame interpolation for single cell tracking. Our cellpose model was trained on 23 semi-automatically labelled images of *U. maydis* and *S. cerevisiae* cells as input (∼ 50 cells/per image). For tracking, micrographs were converted to three-layered (R, G, B) 8-bit depth PNGs before frame interpolation or single-layered 16-bit TIFs before segmentation using the Python package YeastVision. Tracking was performed on hybrid real-interpolated time series, which were downsampled to the original frame rate and used for calculating cell cycle duration, defined as the interval between two budding events. The first detection of a visible bud defined the time point of budding. Budding cells that entered the microfluidics device were excluded from the analysis. Differences between treatments were evaluated assuming each microfluidic device as a biological replicate and using the Kolmogorov-Smirnov test kstest2(), with significance at p < 0.05.

### Coevolution analysis

A dataset of Gpe1 homologs was obtained through a HMMER search^43^ using the sequence of *U. maydis* Gpe1 as bait. Corresponding sequences for all identified Gpe1 homologs were retrieved via the UniProt ID mapping tool (https://www.uniprot.org/id-mapping). These sequences were compiled into a multi-FASTA file and aligned using the MUSCLE alignment tool (https://www.ebi.ac.uk/jdispatcher/msa/muscle?stype=protein). The resulting ‘.clw’ alignment file was used as input for IQ-TREE (https://iqtree.github.io/), which was run locally in a Linux environment. IQ-TREE was executed with default settings, allowing automatic model selection for the dataset prior to calculating the final phylogenetic tree (see original logfile). The resulting ‘.treefile’ output was visualized in the iTOL (https://itol.embl.de/) web application, and the resulting tree was exported as an ‘.svg’ file for figure preparation.

To assess the presence of Pit2 homologs, genomic regions containing Gpe1 homologs were manually examined using Ensembl Fungi (https://fungi.ensembl.org/). When a Pit2 homolog was identified, the corresponding gene names and protein sequences were listed (**Extended Data Table 5**) and used for further alignments using MUSCLE to identify sequence conservation in the consensus motif.

