## Extended Data Figures for "A co-evolved peptide-GPCR system senses host entry to drive fungal infection"

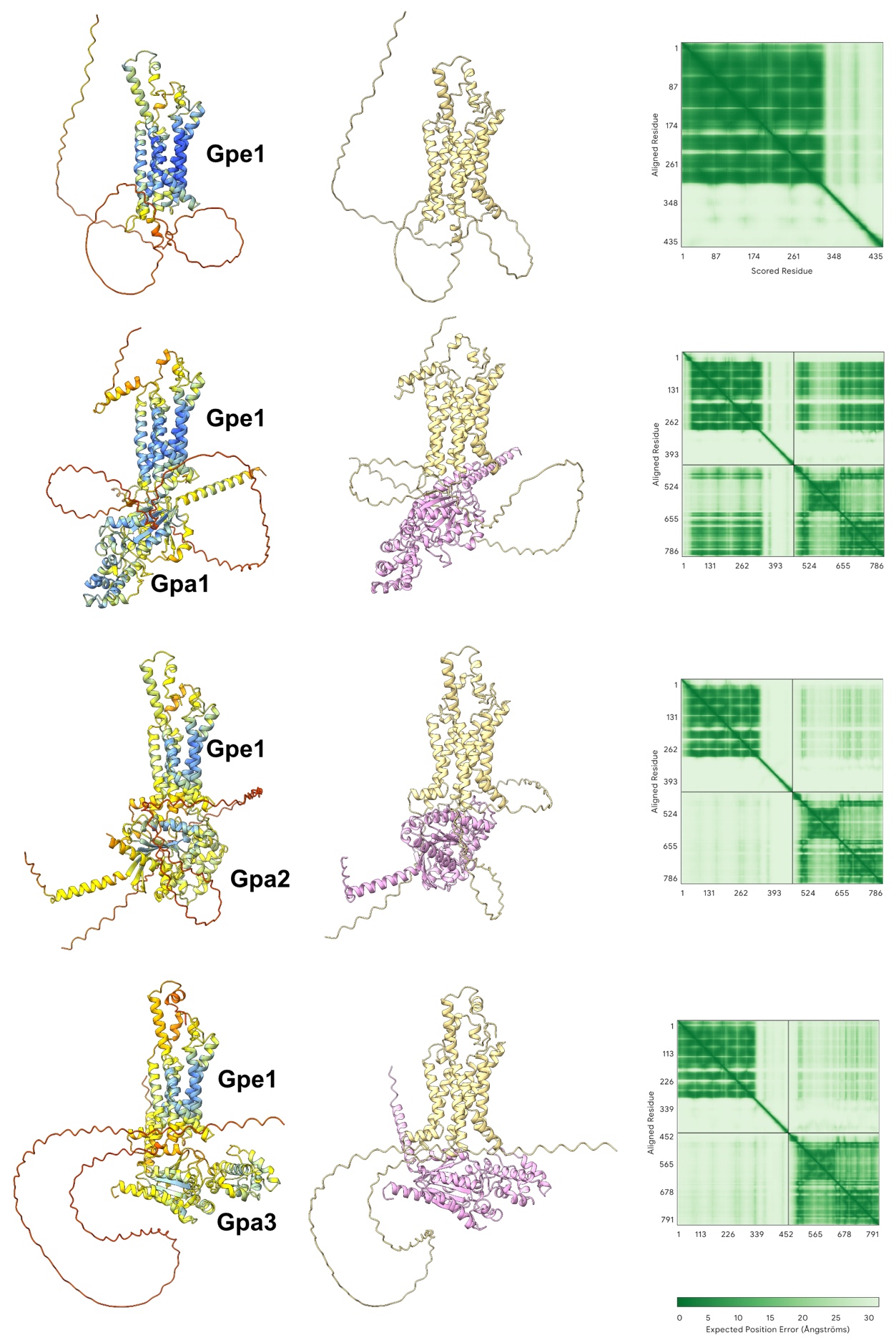

**Extended Data Figure 1.** Structure predictions of Gpe1 alone and in complex with Gpa1, Gpa2 and Gpa3 from *U. maydis*. Shown are the models colored by predicted local distance difference test (pLDDT, left) and molecule (middle). The predicted aligned error (PAE) matrix is shown on the right.

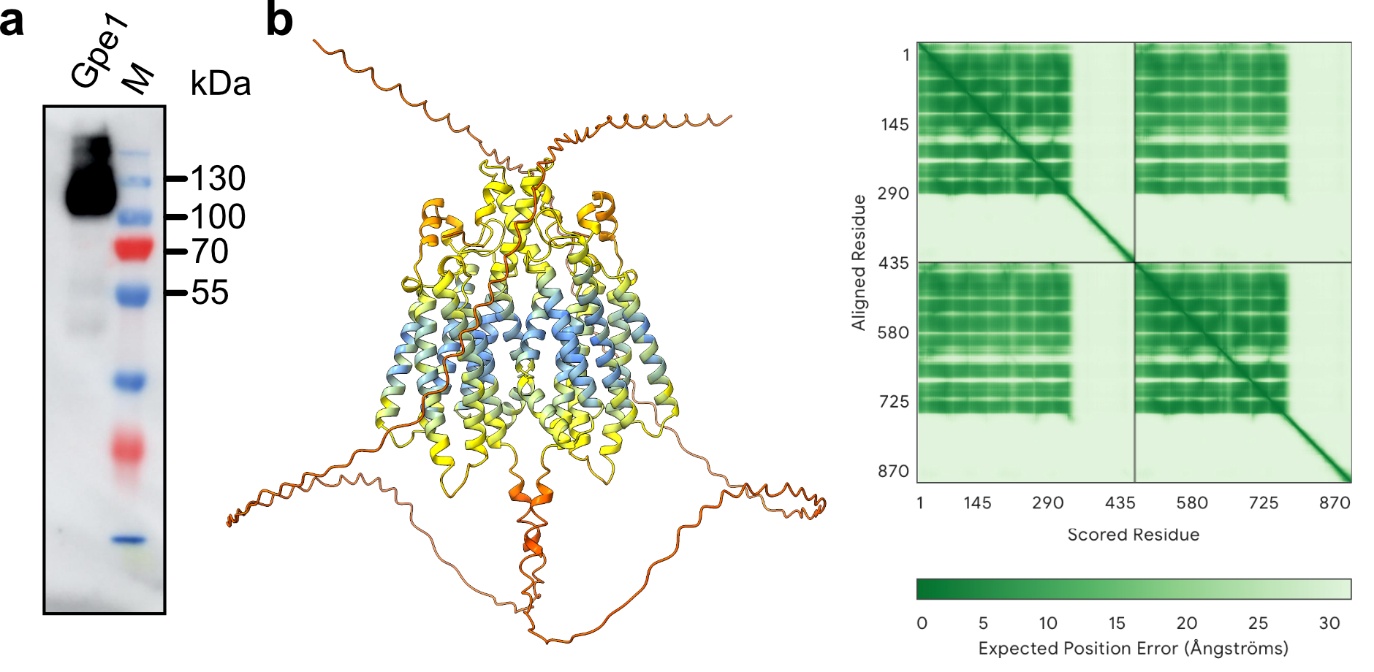

**Extended Data Figure 2.** **Homodimerization of Gpe1 a.** Western blot of Gpe1 expression in *S. cerevisiae.* **b.** Structure prediction of a Gpe1 dimer. Shown is the model colored by predicted local distance difference test (pLDDT, left). The predicted aligned error (PAE) matrix is shown on the right.

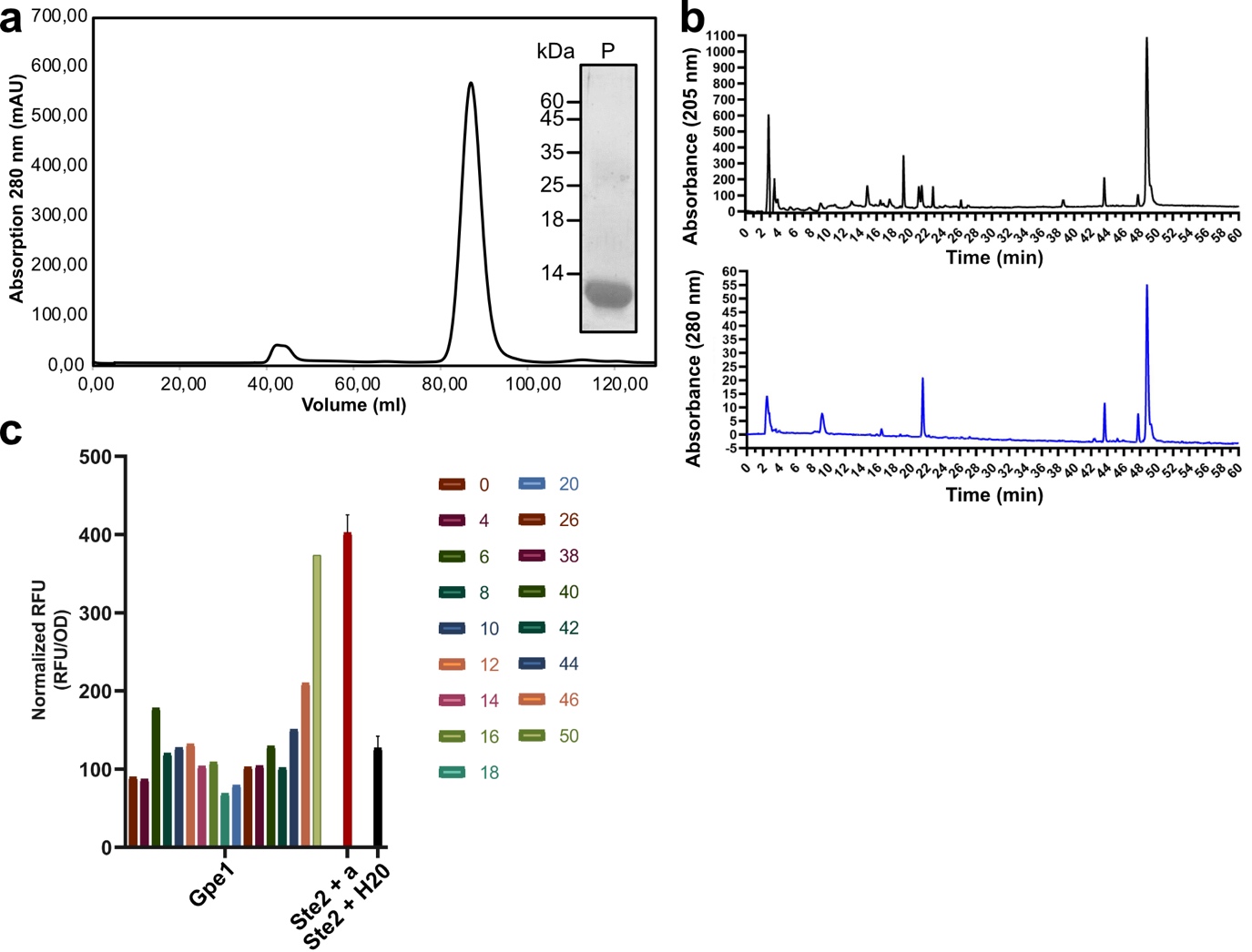

**Extended Data Figure 3.** **Identification of Gpe1-activating Pit2 peptides. a.** Size-exclusion chromatography (SEC) profile of purified Pit2 protein, showing a single monodisperse peak. Inset: Coomassie-stained SDS–PAGE gel of the peak fraction (P). **b.** HPLC analysis of Pit2-derived peptides monitored at 205 nm (top) and 280 nm (bottom), demonstrating the presence of distinct peptide species. **c.** Functional assay comparing Gpe1 receptor activation in response to fractionated peptides. RFU values are normalized to cell density (OD). Gpe1 only responds to fractions 46 and 50. Ste2 activation by the α-factor (red bar), serves as control.

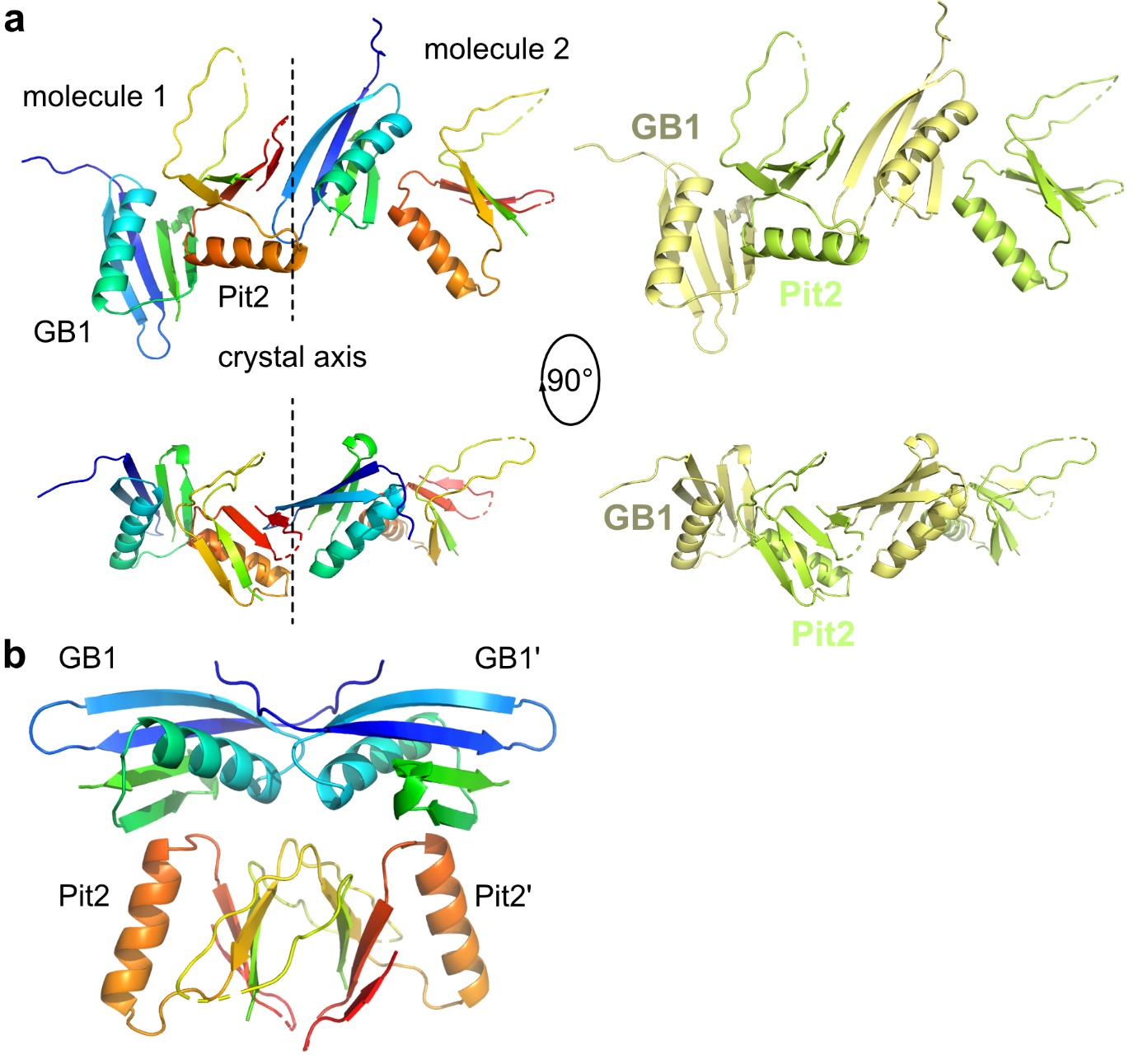

**Extended Data Figure 4.** **Crystallographic analysis of GB1-UmPit2.** **a.** Display of crystallographic symmetry mates of GB1-Pit2 showing that the β-strand 2 of a neighboring GB1 molecule performs a β-completion with β5 of Pit2 stabilizing crystal contacts. **b.** Homodimer of GB1-Pit2 displayed in cartoon style and colored in rainbow from N- to C-terminus.

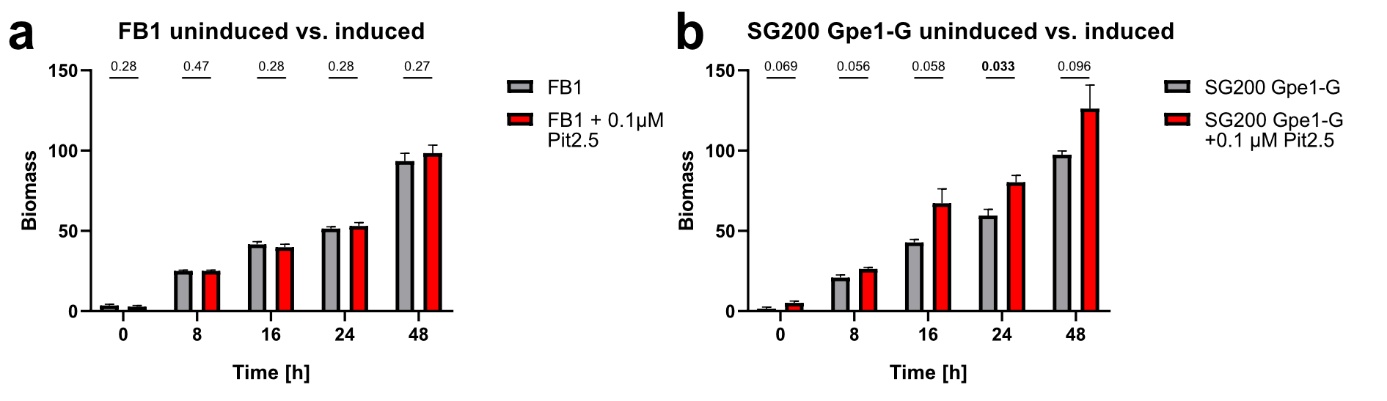
**Extended Data Figure 5.** **Stimulation of Gpe1 by Pit2 increases fungal biomass. a.** *U. maydis* FB1 grown in absence (grey) and presence (red) of 0.1 µM WWFGFTGSL (Pit2.5) peptide. No significant changes between the two conditions are observed. **b.** *U. maydis* SG00 Gpe1-G grown in absence (grey) and presence (red) of 0.1 µM WWFGFTGSL (Pit2.5) peptide. Peptide stimulation increases fungal biomass. P-values obtained from statistical analysis are depicted above each column.

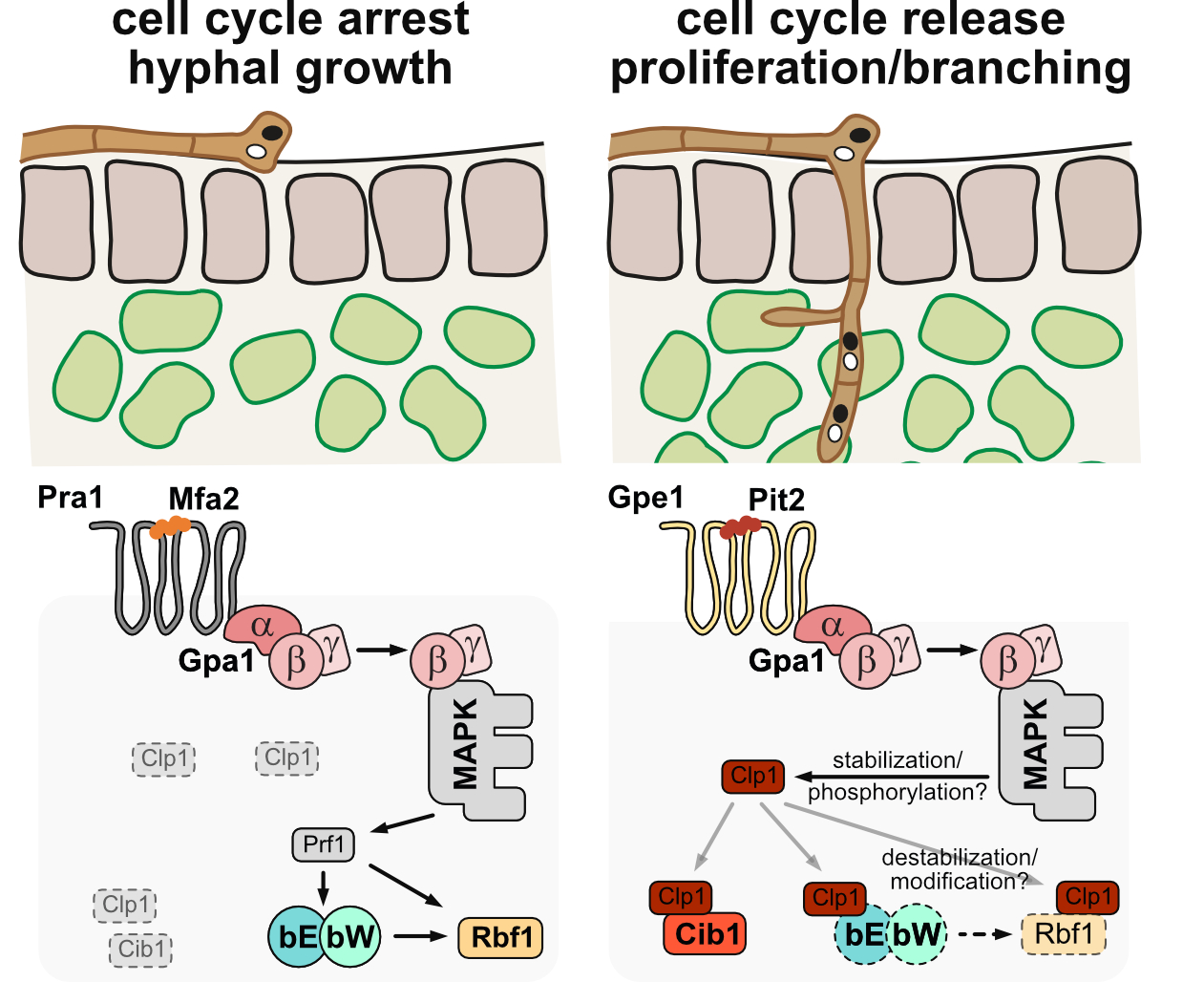

**Extended Data Figure 6. Schematic model of the Gpe1/Pit2 function.** Dashed boxes indicate destabilization/modification of the respective proteins.

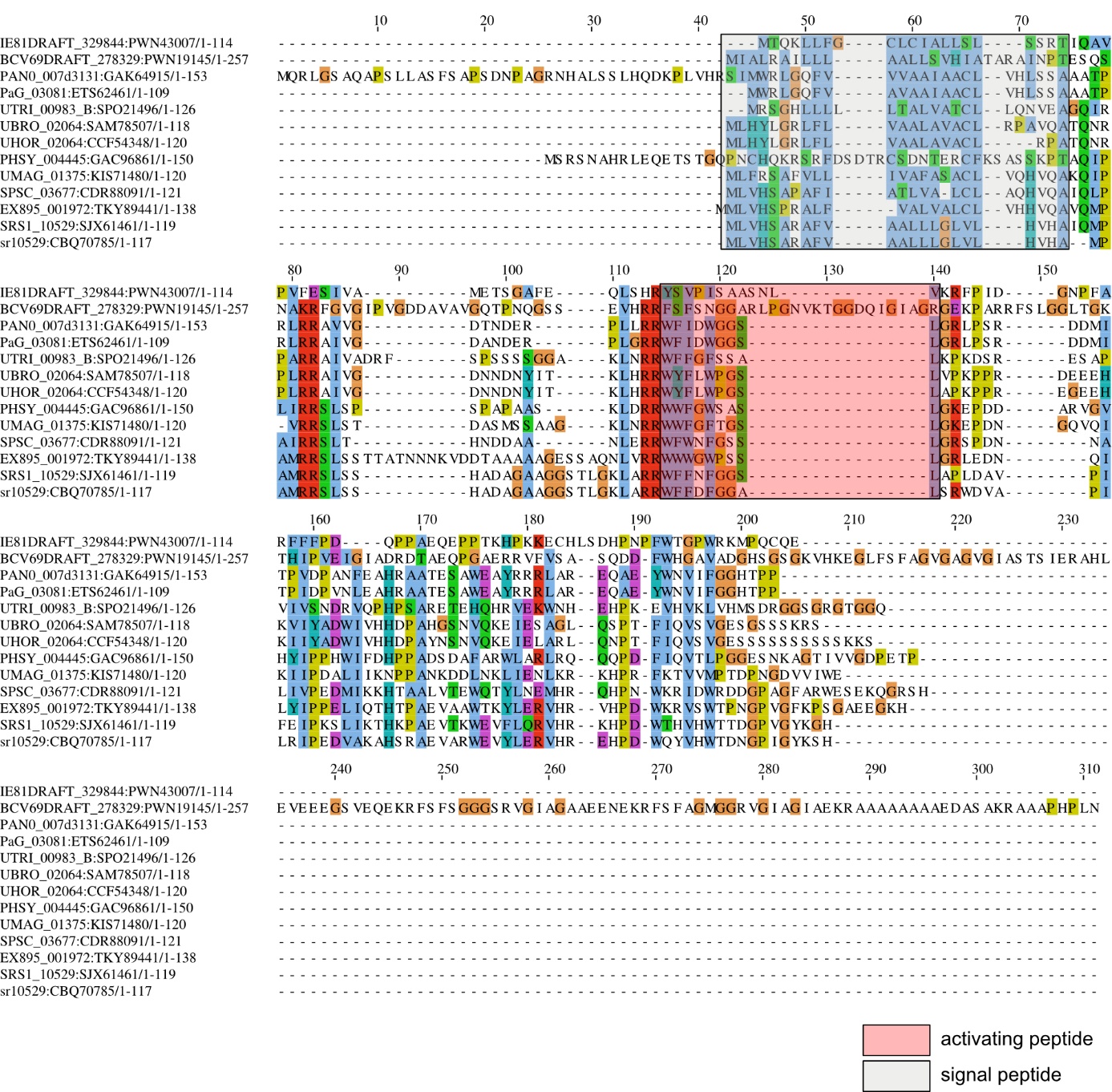

**Extended Data Figure 7.** **Sequence alignment of Pit2 homologs.** Amino acid sequence alignment of Pit2 colored according to the CLUSTAL coloring scheme. The regions containing the signal peptide and the activating Gpe1 ligand are framed in grey and red boxes, respectively.
