## Supplementary Videos for "A co-evolved peptide-GPCR system senses host entry to drive fungal infection": Supplementary_videos.docx

**Video 1.** Representative Pit2 peptide-exposed single cell *U. Maydis* time series in microfluidics. Top right: phase contrast channel with mask contour overlaid in yellow on top of the tracked cell. Top left: GFP fluorescence channel displaying the expression of Gpe1-GFP. Bottom left plot: single cell size quantification as the area in pixels occupied by the tracked cell. Bottom right, single cell mean fluorescence quantification as the average intensity of the pixels occupied by the tracked cell in the GFP channel. Notice the cell divides four times vs three times in the control (Video 2)

**Video 2.** Representative control single cell *U. Maydis* time series not exposed to Pi2 peptides in microfluidics. Top right: phase contrast channel with mask contour overlaid in yellow on top of the tracked cell. Top left: GFP fluorescence channel displaying the expression of Gpe1-GFP. Bottom left plot: single cell size quantification as the area in pixels occupied by the tracked cell. Bottom right, single cell mean fluorescence quantification as the average intensity of the pixels occupied by the tracked cell in the GFP channel. Notice the cell divides three times vs four times in the Pit2-exposed strain (Video 1).
